# Novel quantitative trait loci conferring broad-based resistance to root-knot nematodes in lima bean (*Phaseolus lunatus*)

**DOI:** 10.64898/2026.06.30.735594

**Authors:** Adia M. Tajima, William C. Matthews, Tra Duong, Tran Dang Khanh, Anil Baniya, R. Varma Penmetsa, Travis Parker, Andrew Farmer, Stephanie English, Christine Diepenbrock, Paul Gepts, Philip A. Roberts, Bao-Lam Huynh

**Affiliations:** Department of Nematology, University of California, Riverside, CA 92521; Department of Botany and Plant Sciences, University of California, Riverside CA 92521; Department of Plant Sciences, University of California, Davis, CA 95616; National Center for Genome Resources, Santa Fe, NM 87505

**Author notes:** Corresponding author: Bao-Lam Huynh.

**Keywords:** Lima bean, genetic resistance, Root-knot nematode, *Meloidogyne*, Plant Breeding

## Abstract

Lima bean *(Phaseolus lunatus)* is a broadly adapted, economically important leguminous crop and a susceptible host of root-knot nematodes (*Meloidogyne* spp.; RKN), which are a devastating plant pathogen in agricultural systems worldwide. To date, there have been few studies to elucidate the genetic determinants of RKN resistance in lima beans. Understanding the genetic mechanisms underlying resistance is essential for improving resistance traits and incorporating them into lima bean breeding programs. To assist in marker-assisted selection, we aimed to identify and map quantitative trait loci (QTLs) conferring RKN resistance-related traits. Three recombinant inbred line (RIL) populations were used in this study. Three populations were derived by crossing two RKN-resistant parents with the same RKN-susceptible parent and with each other. All populations were genotyped using genome-wide single-nucleotide polymorphism (SNP) markers. Each population was screened for root galling (RG) and RKN egg reproduction (ER) in response to *M. incognita* and *M. javanica* in greenhouse experiments. Three major QTLs were detected and mapped on chromosome Pl04 (*QRk-pl04.1*), Pl05 (*QRk-pl05.1*) and Pl10 (*QRk-pl10.1*) across populations. Among them, *QRk-pl05.1* and *QRk-pl10.1* affected levels of RG and ER of both RKN species, while *QRk-pl04.1* suppressed root galling and reproduction responses of *M. incognita* but not of *M. javanica*. These chromosomal regions defined by flanking markers will help guide marker-assisted breeding and gene discovery for broad-based RKN resistance in lima beans.

## Introduction

Lima bean *(Phaseolus lunatus)* is a leguminous crop in the Fabaceae family that is valued for its nutrient-dense and edible seeds. Domesticated separately in Mesoamerica and the Central Andes of Ecuador and northern Peru (Gutiérrez-Salgado et al. 1995; Chacón-Sánchez and Martínez-Castillo 2017; Garcia et al. 2021), lima beans have made their mark in the cultural and historical practices of people across the Americas (Marsh n.d.). Today, lima beans are cultivated in over 80 countries and contribute positively to global food security (Traverso et al. 2024). However, lima bean production and the food security it provides may be threatened due to disease susceptibility. One of the most devastating pathogens for lima beans and agriculture worldwide are the root knot nematodes (RKN) (Giordani et al. 2021, Huynh 2016). Among them, *Meloidogyne javanica* and *Meloidogyne incognita* are prevalent in lima bean production regions (Long et al 2014). These nematodes parasitize plant roots by invading the root vascular tissue and inducing root cell differentiation into giant cells that cause root galling. The RKN juveniles live and feed within these giant cells. Although rarely lethal, galling and stress caused by RKN can lead to severe root damage, resulting in reduced water and nutrient uptake, stunted growth, and vegetative chlorosis (Giordani et al. 2021). Yield loss from RKN damage in susceptible crops can range anywhere from 20 - 90% (Garcia et al. 2021). While California’s deep loamy soils are well-suited for lima bean growth, they also foster higher reproduction and penetration rates of RKN in susceptible varieties, resulting in yield losses of up to 70% (Traverso et al. 2024, Long et al 2014, Peterson 2023).

Current pest management strategies to combat the effects of RKN on crops such as lima beans are not highly effective or feasible on a large scale. Although some chemical nematicides are available, they are expensive, and there are significant concerns surrounding their potential health risks. Due to these concerns, some nematicides are banned from use (Giordani et al. 2021, Abad et al. 2008) and a safe alternative to sustainably manage nematode populations is necessary. Unfortunately, crop rotation approaches to reduce pathogenic effects are not highly effective either, as RKN targets a wide variety of crops and the eggs can persist in the soil for multiple years (Giordani et al. 2021). The most promising method of RKN disease and related crop loss management is the use of cultivars with inherent immunity and genetic resistance (Traverso et al. 2024).

The L-136 accession of lima bean (Allard 1954; Helms et al. 2004) was reported to have resistance to *Meloidogyne incognita* (reduced galling and reproduction) and *M. javanica* (reduced galling, but normal reproduction), in a previous study by Roberts et al. (2008). In that study, quantitative inheritance analysis of a Henderson x L-136 recombinant inbred line (RIL) population indicated that root galling and reproduction were controlled by different unidentified genes based on phenotypic data from greenhouse and field tests. However, genomic regions and markers for these genes have yet to be reported. The PI 256874 accession of lima bean (PI 256874 GRIN-Global 2025), known by G code G25467, was selected as a second RKN resistant parent used to develop recombinant inbred lines. PI 256874 exhibits indeterminate growth and is photoperiod sensitive.

L-136 and PI 256874 were selected as two resistant parents to cross with the commercial lima bean cultivar Henderson and with each other to develop three F_2:9_ RIL populations: Henderson x L-136, Henderson x PI 256874, and PI 25684 x L-136. In this study, the three populations were genotyped using Restriction site-Associated DNA sequencing (RAD-Seq), resulting in 739, 2278, and 1759 genome-wide SNP markers, respectively. The populations were phenotypically scored for galling and reproduction response to *M. incognita* and *M. javanica* in greenhouse pot trials. We report on quantitative trait loci (QTLs) significantly associated with RKN resistance in the three lima bean RIL populations. Flanking markers for these QTL regions can help lead future gene discovery endeavors and marker-assisted backcrossing for the development of RKN-resistant lima bean cultivars.

## Materials and Methods

### Mapping Populations

Three segregating lima bean populations were used in genetic mapping. The RIL01 population included 119 F_9_ RILs derived from a cross between the susceptible cultivar Henderson as a female parent and the resistant accession L-136 as male parent (Roberts et al. 2008). The RIL02 population included 107 F_9_ RILs derived from a cross between Henderson (female) and another resistant accession PI 256874 (male) (Roberts et al. 1998). The RIL03 population includes 113 F_9_ RILs derived from a cross between the two resistant parents PI 256874 (female) and L-136 (male). Henderson is a commercial variety of lima beans that are highly susceptible to both *Meloidogyne* species *M. incognita* and *M. javanica*.

### Nematode Isolates

Two RKN isolates were used to inoculate and phenotype plants for galling and reproduction response. The *Meloidogyne incognita* isolate “Beltran” was originally isolated from a lima bean field in the San Joaquin Valley in California. The *Meloidogyne javanica* isolate “811” was originally isolated from a cowpea field in Southern California. Both “Beltran” and “811” isolates were maintained on greenhouse grown tomato plants susceptible to RKN stress (Thomason and McKinney 1960). Nematode eggs were extracted from susceptible tomato roots and used to inoculate greenhouse grown lima bean RILs and parents for resistance phenotyping.

### Resistance Phenotyping

The three RIL populations were previously developed and phenotypically scored for their response to RKN stress at UCR in the early 2000s. Phenotypic data of the RIL01 population were reported in Roberts et al. (2008). All RIL populations were separately screened in greenhouse pot trials for resistance to *M. incognita* and *M. javanica* galling and reproduction response. Individual plants were grown in plastic pots with steam-sterilized 80:20 sand: peat moss mix. Plants were inoculated with 50,000 eggs in 10 mL of water of *M. javanica* and *M. incognita* respectively after the first set of trifoliate leaves emerged, approximately 3 weeks after germination. The RKN hatching and infiltration of root systems began within several days of initial inoculation of greenhouse pot tests. Symptoms of RKN infestation, including galling of root systems and development of nematode egg masses visible on the root surface, were used for phenotyping. Roots of RIL lines and parents were washed clean after 2 months of growth and scored for root galling. The RIL01 population was scored differently from the RIL02 and RIL03 populations as specified below.

The RIL01 population’s root systems screened in greenhouse pot trials were scored for *M. incognita* and *M. javanica* galling and reproduction (Roberts et al. 2008) using a rating scale from 0 (resistant, no gall/egg mass) to 4 (galls/egg masses present across the root system). The egg mass production was measured by staining egg masses with an erioglaucine solution (Omwega et al 1988). Four replicates (pots) of each line were measured, and the mean score was used in QTL analysis.

The RIL02 and RIL03 populations were visually scored for galling on a scale of 1-10 (Bridge and Page 1980). Egg production was scored by extracting egg masses from root systems using a 0.5% NaOCl solution and vigorous shaking in an industrial paint shaker. Eggs per root system (ERS) for the RIL02 and RIL03 populations were calculated by counting extracted eggs using a tabletop Zeiss light microscope and cell counting slides. The egg: root mass ratio trait was determined by weighing root systems and dividing the ERS value by their root weight, to provide an eggs/gram root (EGR) estimate. The ERS and EGR scores were log-adjusted by calculating the log_10_(n+1) score and using those values in phenotype histograms and QTL analysis.

### Marker genotyping

Leaf tissue of all three RIL populations was collected at UC Riverside from greenhouse-grown plants. DNA of the RIL01 population was extracted using the Qiagen DNEasy Plant Minikit according to the manufacturer’s protocol. Leaf tissue of individual RIL02 and RIL03 plants were prepared in a two-step protocol, with an initial DNA extraction done using the Hu and Lagarias method (Hu and Lagarias 2020) that was followed by a column purification with a Zymo Research ZR-96 DNA clean up kit according to manufacturer’s protocol. Purified DNA for all three RILs were quantified by PicoGreen fluorometry, normalized to equal concentrations and SNP genotyped at the UC Davis Genome Center’s core genotyping facility using reduced representation Restriction site-Associated DNA sequencing (RAD-Seq) using the single enzyme Nla-III whose isoschizomer CviAII has been previously documented to provide genomewide coverage in lima bean (Ariani et al 2016). Adapters were trimmed from reads using cutadapt (Martin 2011) and subsequently aligned to the reference genome with Minimap2 (Li 2018). Sample variants were identified using GATK HaplotypeCaller (McKenna et al 2010). The R package mpr.genotyping (adf-ncgr 2025) was used to reduce noise in genotypes by minimizing the number of recombination events locally in each sample. Markers with minor allele frequencies greater than 0.2 were retained for linkage analysis.

### Genetic mapping

Linkage analysis and QTL mapping were performed using the QTL IciMapping 4.2 software (Meng et al., 2025). Markers that were inherited together without recombination were grouped into bins, from which one marker per bin was used in map construction and QTL analysis. Linkage maps were constructed using the Kosambi function, with reference to the SNP physical locations from the UC92 x UC Haskell reference genome (Garcia et al. 2021). A QTL analysis was performed using the Inclusive Composite Interval Mapping (ICIM) method (Wang 2009) run with the QTL IciMapping software. In brief, the ICIM involved three consecutive steps: (1) Single marker analysis was used to select significant markers (*P* < 0.001) associated with phenotypes, (2) phenotypic values were adjusted for the selected markers except for the two markers flanking the current mapping interval, and (3) the adjusted phenotypic values were used in composite interval mapping, which involves testing QTL additive effect between QTLs.

## Results

### Phenotypic variation

Bimodal distributions were observed for root galling response measured in the RIL01 population assayed with *M. incognita* and *M. javanica*, with the resistant parent L-136 showing a lower galling than the susceptible parent Henderson (Figure 1a,c). The *M. incognita* egg masses measured in the RIL01 population also show a bimodal distribution (Figure 1b) but were not significantly correlated (*r* = 0.135; *P* > 0.05) with the root galling response. The *M. javanica* egg masses were produced in all RILs and parents of this population and were evenly distributed among RIL lines.

**Fig. 1.**
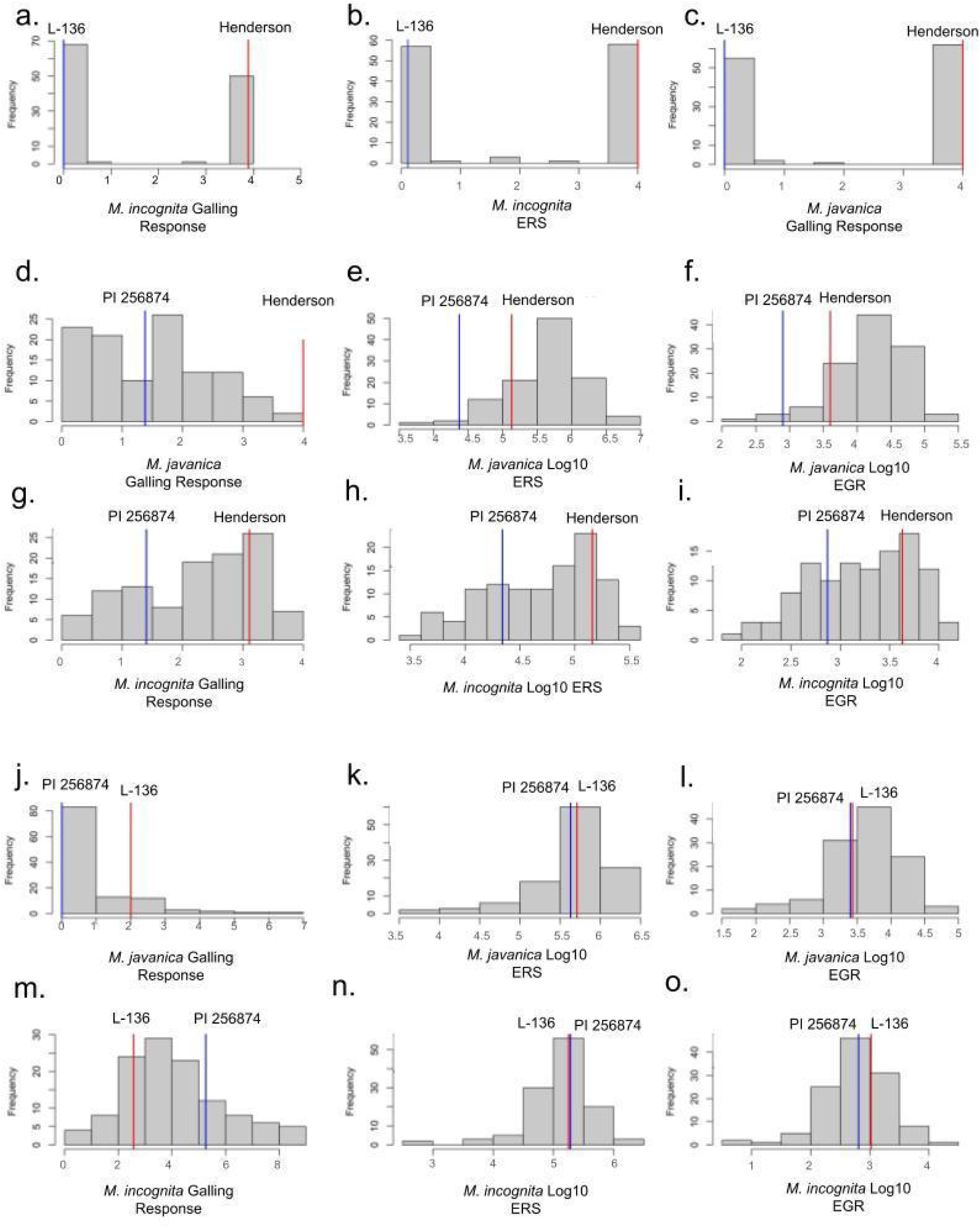
Variation in root-knot nematode galling and reproduction response (ERS or EGR) of the RIL01, RIL02 and RIL03 populations. ERS: Eggs per root system, EGR: number of eggs/g of roots

The galling phenotype in the RIL02 population showed a bimodal distribution in response to *M. incognita* and *M. javanica* (Figure 1d,g). For *M. javanica* ERS and EGR (Fig. 1e,f), the distribution of phenotypes was skewed towards the Henderson phenotype, with PI 256874 and Henderson both having a relatively lower reproduction response compared to the average distribution of the population. There was a significant correlation between ERS and EGR (*r* = 0.663, *P* < 0.001) in *M. javanica*, however, their correlation with root galling response was not significant (*r* = 0.323 and 0.158, respectively; *P* < 0.05). For *M. incognita* ERS and EGR (Fig. 1h,i), the population was more evenly distributed with the PI 256874 line showing less galling and reproduction response overall compared to Henderson. There was a significant correlation between ERS and galling response in *M. incognita* (*r* = 0.62, *P* < 0.001), as well as ERS and EGR (*r* = 0.737, *P* < 0.001).

The RIL03 population showed wide variation in reproductive and total EGR responses to *M. incognita* and *M. javanica,* while both parents had similar values (Figure 1k,l,n,o). The reproduction data were generally skewed towards the higher value whereas the parental lines were centered. In the RIL03 population, there was a negligible correlation between *M. javanica* reproduction and galling, whereas there was a strong correlation between *M. incognita* galling and reproduction response (*r* = 0.717, *P* < 0.001). Galling data showed that L-136 was more susceptible to *M. javanica* galling, while PI 256874 exhibited higher susceptibility to *M. incognita* galling (Figure 1j,m). Galling data for RIL03 did not follow a bimodal distribution like those seen in the RIL01 and RIL02 populations. Significant correlations between *M. incognita* reproduction and galling response were observed in both the RIL02 and RIL03 populations (*r* = 0.62 and 0.72, respectively; *P* < 0.001).

### QTL Identification

Three significant QTL peaks were identified on chromosomes Pl04, Pl05, and Pl10 in the RIL01 population (Figure 2a,b,c). A major QTL on Pl04 corresponded to the *M. incognita* galling response, explaining some 90% of total phenotypic variance (PVE) (LOD = 125.85, PVE = 89.59%). Another major QTL region mapped to Pl05 and was associated with *M. incognita* egg mass and reproduction response (LOD = 78.44, PVE = 92.04%). The *M. javanica* galling response phenotype was associated with a major-effect QTL region on chromosome Pl10 (LOD = 110.1, PVE = 91.99%) (Figure 2c). The favorable (low galling and reproduction) allele was contributed by the resistant parent L-136.

**Fig. 2.**
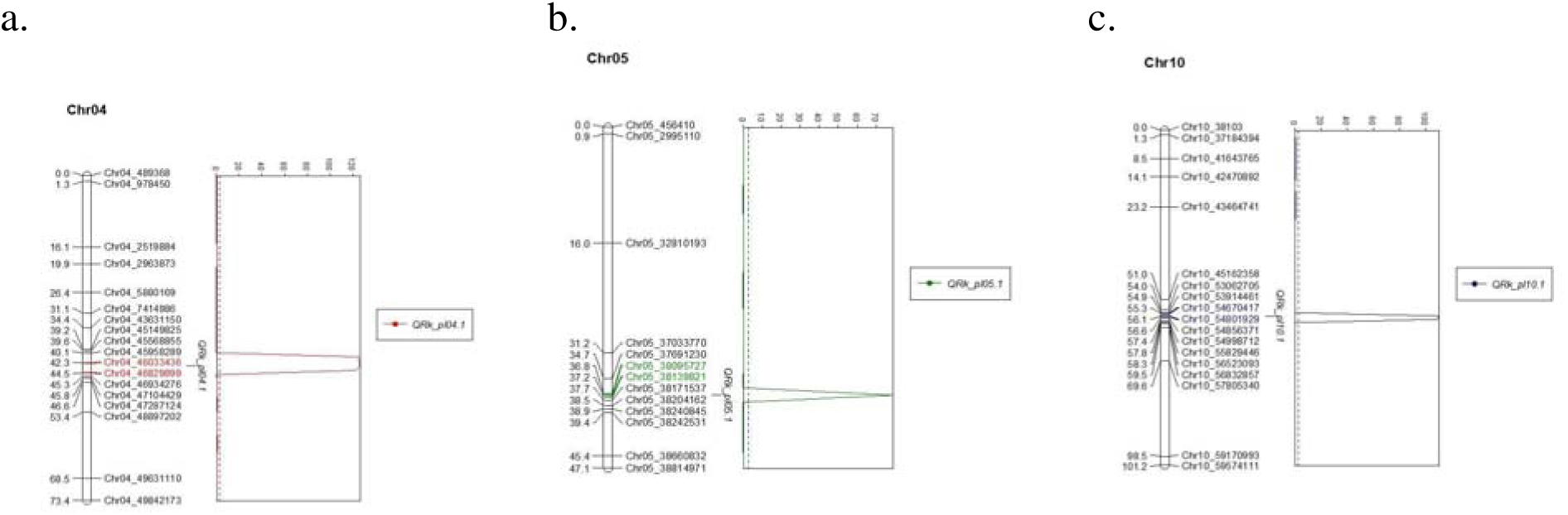
Chromosomal regions associated with (a) *M. incognita* galling, (b) *M. incognita* reproduction, and (c) *M. javanica* galling in the RIL01 (Henderson x L-136) population. Markers flanking the QTL peak are colorized. Chr04, Chr05, Chr10: Chromosomes Pl04, Pl05, and Pl10, respectively. Numbers on the left of each marker represent chromosomal location in centimorgans.

In the RIL02 population, a major QTL region was identified on chromosome Pl05 in significant association with *M. incognita* and *M. javanica* galling, reproduction, and EGR traits (Figure 3a,b). The LOD and PVE values varied within these traits, with the highest LOD and PVE scores being associated with *M. incognita* ERS and EGR (LOD = 19.73 and 20.58, PVE = 54.8% and 60.1%, respectively), followed by *M. javanica* ERS and EGR (LOD = 8.3 and 8.1, PVE = 34.12% and 33.44%, respectively). The QTL on Pl05, associated with *M. incognita* galling, ERS, and EGR and *M. javanica* galling overlapped with the QTL on Pl05 associated with *M. incognita* egg mass from the RIL01 population. The favorable (low galling and reproduction) allele was contributed by the resistant parent PI 256874.

**Fig. 3.**
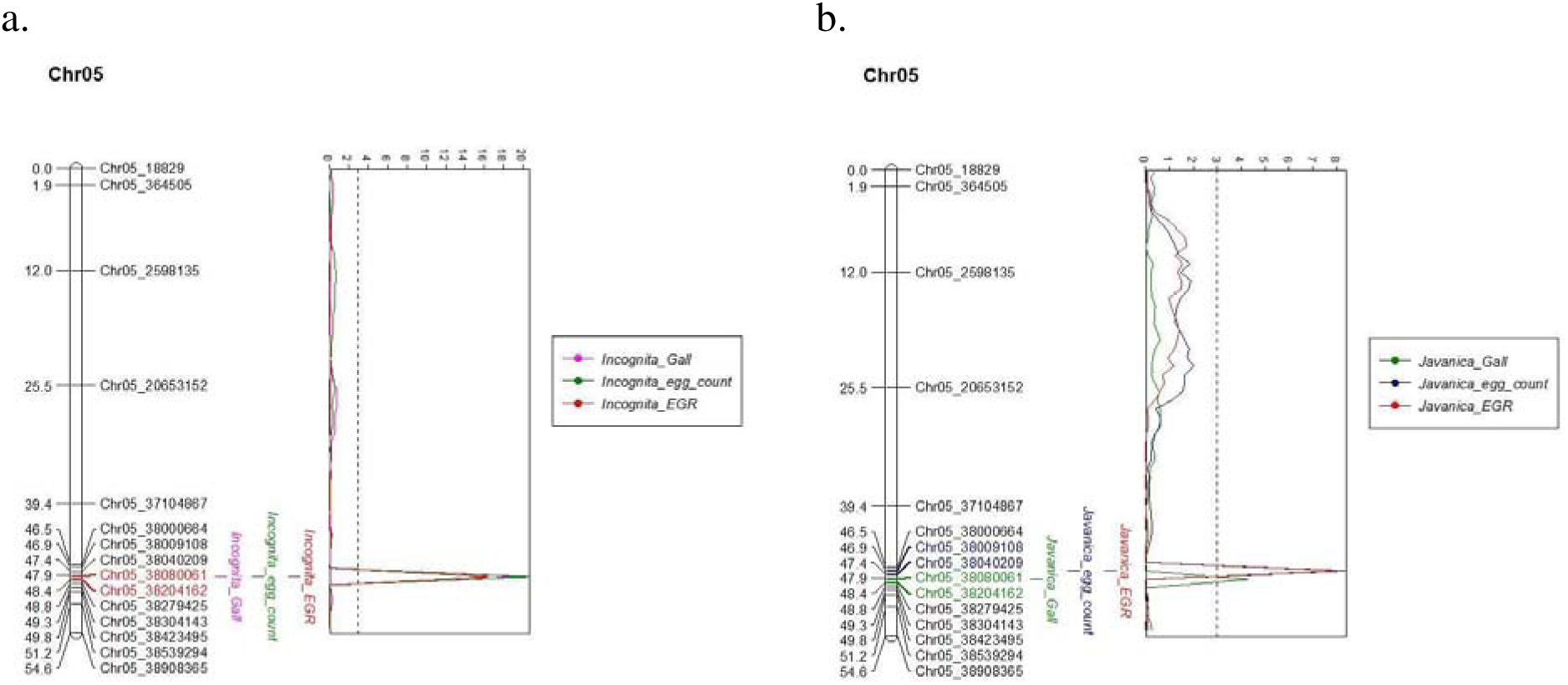
Chromosomal regions associated with galling, egg count and egg count: root ratio response to (a.) *Meloidogyne incognita* and (b.) *Meloidogyne javanica* in the RIL02 (Henderson x PI 256874) population. Markers flanking the QTL peak are colorized. Chr05: Chromosome Pl05; ERS: Eggs per root system, EGR: eggs/g of roots.

The RIL03 population exhibited a major QTL on chromosome Pl04 in significant association with *M. incognita* galling (LOD = 22.8; PVE = 58.9%), ERS (LOD = 12; PVE = 31.4%) and EGR (LOD = 221.8; PVE = 17.4%), with the favorable (low galling and reproduction) allele being contributed by the parent L-136. (Figure 4a; Table 1). There was also a minor QTL on Pl05 associated with *M. javanica* and *M. incognita* EGR (LOD = 4.4 and 5.8; PVE = 3.1% and 3.2%, respectively), which overlaps with the QTLs identified in the RIL01 and RIL02 populations (Figure 4b). Another QTL mapped to Pl10 in significant association with *M. incognita* EGR (LOD = 31.9, PVE = 33.2%), with the favorable (low galling and reproduction) allele being contributed by the parent PI 256874 (Figure 4c; Table 1). Finally, Pl08 harbored two QTLs associated with *M. javanica* EGR ( LOD=12.4 and 20.4, PVE=10.5 and 21.1%), with the favorable (low galling and reproduction) allele contributed by parents L-136 and PI 256874, respectively (Figure 4d and Table 1).

**Fig. 4.**
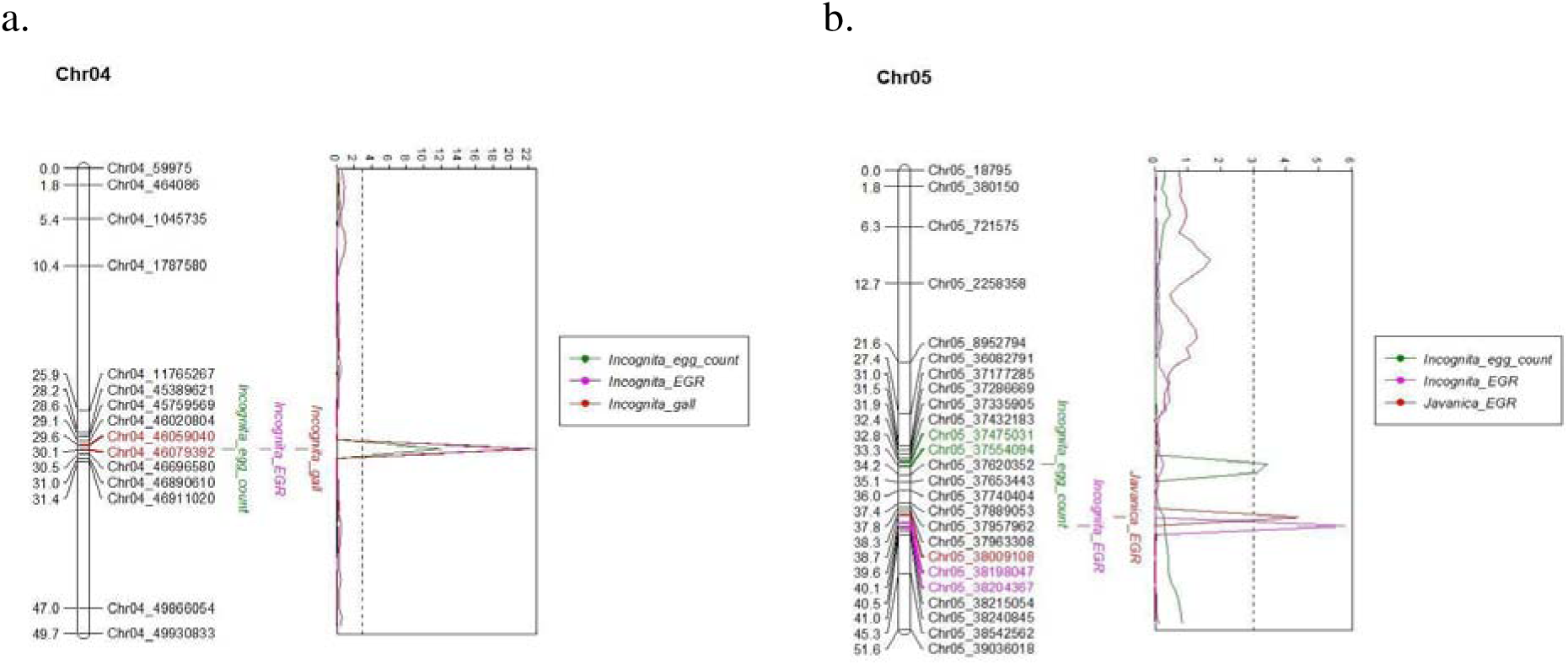
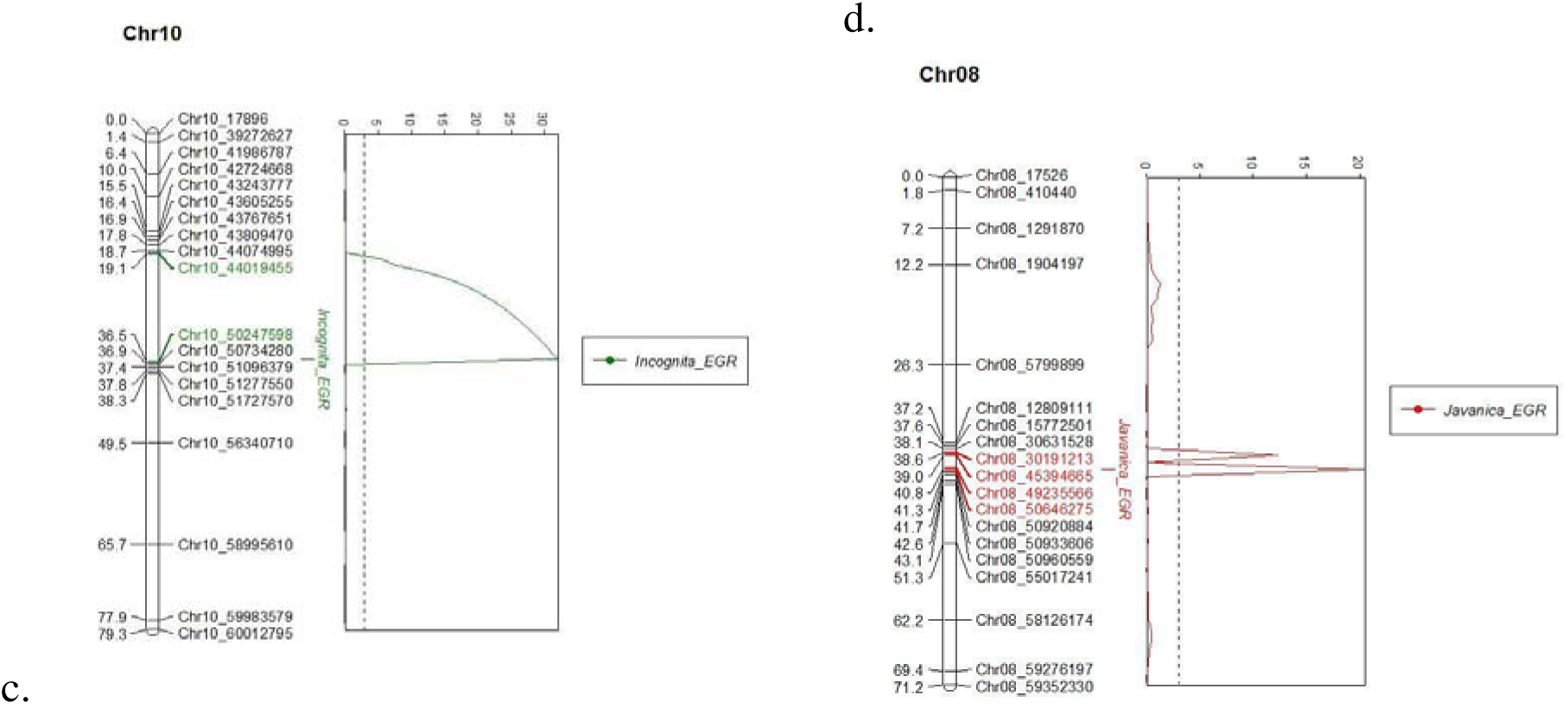
Chromosomal regions associated with root-knot nematode resistance in the RIL03 (PI 256874 x L-136) population. (a.) depicts LOD peaks associated with *M. incognita* egg count (green), *M. incognita* EGR (pink) and *M. incognita* galling (red) on chromosome 4. (b.) depicts LOD peaks associated with *M. incognita* ERS (green), *M. incognita* EGR (pink) and *M. javanica* EGR (red) on chromosome 5. (c.) depicts LOD peaks associated with *M. incognita* EGR on chromosome 10. (d.) depicts LOD peaks associated with *M. javanica* EGR on chromosome 8.Markers flanking the QTL peak are colorized. Chr04, Chr05, Chr08, Chr10: Chromosomes Pl04, Pl05, Pl08, and Pl10, respectively. ERS: Eggs per root system, EGR: eggs/g of roots

**Table 1.**
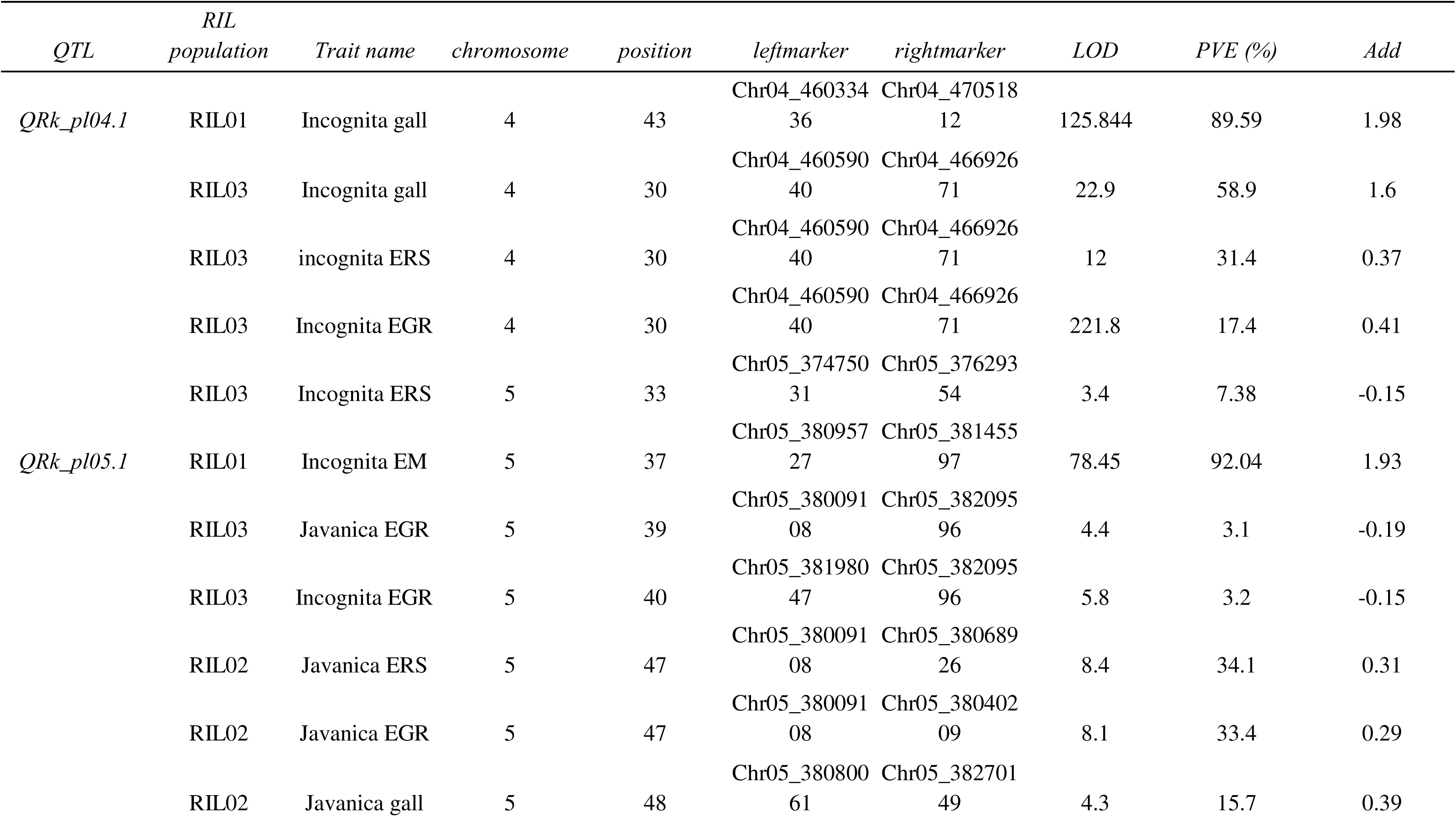
Quantitative trait loci associated with root galling and reproduction response to different root-knot nematode species assessed in three RIL populations Henderson x L-136 (RIL01), Henderson x PI 256874 (RIL02), and PI 256874 x L-136 (RIL03).

## Discussion

The bimodal distribution of the root galling and reproduction phenotypes in the RIL01 and RIL02 populations (Figure 1a, b, c, d, g) supports the presence of a major QTL in each RIL population. The QTL peak on chromosome Pl04 associated with *M. incognita* galling and reproduction response in the RIL03 population spans approximately 633 kb, which falls within the 1000-kb QTL (∼2 cM) on the same chromosome associated with *M. incognita* galling in the RIL01 population (Figures 2 and 4). Hereafter, this QTL region is referred to as *QRk-pl04.1*. There was also overlap among QTLs on chromosome Pl05 mapped in different RIL populations. These include the QTL for *M. incognita* egg mass in the RIL01 population (Figure 2), the QTL for *M. incognita* and *M. javanica* galling and reproduction in the RIL02 population (Figure 3), and the QTL for *M. javanica* and *M. incognita* EGR in the RIL03 population (Figure 4). Hereafter, this QTL region is referred to as *QRk-pl05.1*, which spans around 200 kb based on consensus flanking markers (Chr05_38009108 and Chr05_38209596) across the three overlapping QTL. The other major QTL region on Pl10, hereafter *QRk-pl10.1*, was associated with *M. javanica* galling in the RIL01 population (Figure 2) and *M. incognita* EGR in both the RIL02 and RIL03 populations (Figures 3 and 4), spanning around 5,629 kb based on consensus flanking markers (Chr10_45103536 and Chr10_50732730) across two overlapping QTL. The significant parental effects at *QRk-pl04.1, QRk-pl05.1,* and *QRk-pl10.1* in the RIL03 population indicate that these QTLs are multi-allelic (i.e., L-136 and PI 256874 may carry different resistance alleles at each QTL) and/or are subject to epistatic interactions involving the respective genetic backgrounds of the resistance sources. The presence of common QTLs across populations confirms their conserved effects that could be valuable for marker-assisted breeding. Additional minor QTL peaks detected on chromosomes Pl01, Pl02, Pl03, Pl08, and Pl09 in either the RIL02 or RIL03 populations (Supplemental Figure 1) indicate the presence of minor resistance genes and their possible interactions.

A previous quantitative genetic analysis by Roberts et al. (2008) suggested the presence of three nuclear genes with independent control in the RIL01 cross of Henderson x L-136. These genes are *Mir-1*, a recessive gene that controls *M. incognita* egg production, *Mig-1*, a dominant gene that controls *M. incognita* galling, and *Mjg-1*, a dominant gene that controls *M. javanica* galling. Based on F_1_ data (Supplementary Table 3), the L-136 resistance to galling by *M. incognita* and *M. javanica* is dominant in both cases, while the L-136 resistance to *M. incognita* reproduction is recessive. The QTLs associated with the RIL01 population correspond phenotypically to these previously described genes. *QRk-pl04.1*, associated with *M. incognita* galling response, is consistent with the dominant *Mig-1* gene. *QRk-pl05.1*, associated with *M. incognita* egg mass and reproduction response, corresponds to the recessive *Mir-1* gene. Finally, *QRk-pl10.1* is associated with *M. javanica* galling and is consistent with the dominant *Mjg-1* gene. The QTLs identified in this study offer a genomic localization for these previously described genes and confirm their independent genetic control.

The identification of multiple QTLs for RKN resistance in lima beans is also consistent with previous reports on the multigenic control of RKN resistance in other crops. For example, in common bean (*P vulgaris*), the minimum number of genes involved in RKN resistance was around 2-3 (Pesqueira 2025, Orsi 2025). The concentration of QTLs on chromosome Pl05 in the RIL02 population indicates that a gene, or more likely a complex or cluster of genes, is involved in much of the resistance to *M. incognita* and *M. javanica* in the PI 256874 line. For example, a single gene or complex of linked genes acting as a single gene on Pl05 likely may explain the prior observation that 13 lines exhibited resistance to *M. incognita* galling, *M. incognita* reproduction (low egg counts), as well as somewhat lower *M. javanica* galling, but no resistance to *M. javanica* reproduction in the phenotypic Class 4 (Supplementary Table 1). Perhaps accordingly, the QTL (flanked by Chr05_38009108 and Chr05_38068926) in the RIL02 population associated with resistance to *M. javanica* reproduction did not overlap with the resistance to *M. incognita* (galling and reproduction) and the *M. javanica* galling QTL (flanked by Chr05_38080061 and Chr05_38270149), indicating a possible separate but linked gene control of the *M. javanica* reproduction response. The 17 lines observed in phenotypic Class 3 (Supplementary Table 1) with low *M. javanica* galling but weakened resistance to *M. javanica* reproduction is likely due to the above QTL (flanked by Chr05_38080061 and Chr05_38270149) combined with the resistance associated with the QTL on chromosome Pl09. The *M. incognita* galling trait exhibited a negative additive effect for one of the QTL peaks (Pl03) with which the trait was associated with, indicating PI 256874 as contributing to the unfavorable, or high value, allele. For the rest of the trait-associated peaks, the positive additive effect suggests that Henderson contributes to the high-value allele. This suggests that PI 256874 contributes a majority of the resistance seen in the RIL02 population, and that gene(s) on Pl05 are responsible for most of the observed resistance.

The RIL03 population is somewhat more complex in differentiating between the resistance effects of each parent line. Based on additive effects, the resistance towards *M. incognita* and *M. javanica* was contributed by both L-136 and PI 256874 parents. The *QRk-pl04.1* locus was associated with resistance contributed by the L-136 parent, while the *QRk-pl05.1* and *QRk-pl10.1* loci were associated with resistance contributed by the PI 256874 parent. Interestingly, L-136 and PI 256874 have different geographic origins; L-136 originated from Puerto Rico (of Mesoamerican origin) and PI 256874 originated from Peru (of Andean origin) (Allard 1954, Helms 2004, PI 256874 GRIN-Global 2025). The small and large seed size of L-136 and PI 256874, respectively, confirms their difference in place of origin. This difference raises the possibility that the observed QTLs may not be allelic in nature.

Supplementary Table 2 demonstrates the galling, egg count and egg: root ratio scores associated with lines possessing the resistance QTL haplotypes on chromosomes Pl04, Pl05 and Pl10. For example, lines that possess the L-136 haplotype at *QRk-pl04.1* and the PI 256874 haplotype *QRk-pl05.1* and *QRk-pl10.1* on average had enhanced resistance to *M. incognita* and *M. javanica* (lines 47, 63, 70, 72, 81, 83). There are also examples of lines that do not carry any alleles of the resistant parent at each of the loci (lines 16, 71, 87). In general, lines with resistance QTL haplotypes had lower galling and egg production scores than those with the susceptible QTL haplotype. The RIL03 population also had lines with different combinations of genes resulting in highly resistant lines (more resistant than either resistant parent, e.g., lines 47 & 81, resistance to *M. incognita* reproduction), as well as lines that have lost all resistance (susceptible recombinants, e.g., line 71, no resistance to javanica reproduction). These lines demonstrate the transgressive segregation resulting from the presence of multiple resistance QTLs in the RIL03 population.

There are multiple avenues for further analysis based on these QTL discoveries. Backcrossing analyses can be performed in order to further refine the genic region underlying galling resistance in L-136, as well as to investigate the recessive resistance to galling and reproduction observed in PI 256874. RNA-sequencing is another important avenue to narrow down candidate genes within the QTL regions, as reported for cowpea (Santos et al. 2018) and common bean (Orsi et al. 2025). A potential future experiment could be to select one of the RIL populations to analyze gene expression timelines using RNA sequencing after exposure to nematode stress.

Another future avenue could be to utilize transgressively resistant lines that carry a full complement of the resistance traits from the RIL03 population for marker-assisted selection in lima bean breeding. Overall, our report has helped to elucidate major QTLs associated with RKN resistance, paving the way for marker-assisted backcrossing or further candidate resistance gene characterization.

## Conclusion

This study has identified QTLs of significance across three RIL populations that can be used for marker-assisted breeding of resistance to root-knot nematodes *M. javanica* and *M. incognita* in lima beans. By integrating genotypic data with phenotypic analyses of root galling and nematode reproduction response, we were able to confirm that the *M. incognita-*induced galling and reproduction response, and *M. javanica* galling in the RIL01 population are under separate genetic control. Analyses of the RIL02 and RIL03 populations identified QTLs on chromosomes Pl04, Pl05, and Pl1010 associated with RKN resistance in all three populations. Analyses of F_1_ populations established that resistance to galling in L-136 is dominant, while resistance to reproduction in L-136 and resistance to galling and reproduction in PI 256874 is recessive. This is important for marker-assisted breeding to select for recessive resistance alleles, which is a challenge when using phenotypic selection alone. Further research should focus on refining the QTL regions of interest through backcrossing and gene expression experiments to identify genes in resistant parental lines and to subsequently develop gene-based molecular markers for use in resistance breeding.

## Acknowledgements

This work was supported by grant No. 2022-51181-38323 (to PD Paul Gepts) of the Specialty Crop Research Initiative (SCRI) Program at the United States Department of Agriculture (USDA) and AES Hatch Project (to Philip A. Roberts). The authors wish to thank Peggy Mauk and team at the Agricultural Operations facility on UC Riverside campus for cooperation and trial management. The authors also would like to thank Sarah Dohle for information regarding parental accessions.

## Statements and Declarations

### Author Contributions

P.A.R. and W.C.M. conceived the experiment. W.C.M. made the crosses, developed the RIL lines and collected data. T.D., V.P., A.F., and S.E. contributed to genotyping efforts. B.L.H. constructed the genetic maps and guided the genetic discovery effort. A.M.T. performed data analysis, QTL mapping, manuscript writing, and editing. B.L.H., P.A.R., W.C.M., C.D. and P.G. contributed to manuscript editing. P.A.R. and P.G. obtained funding. All authors read and approved the final manuscript.

### Data Availability

The datasets generated and/or analysed during the current study are available from the corresponding author upon reasonable requests. Datasets are also accessible via Dryad: https://doi.org/10.5061/dryad.79cnp5jc5

**Table 1.**
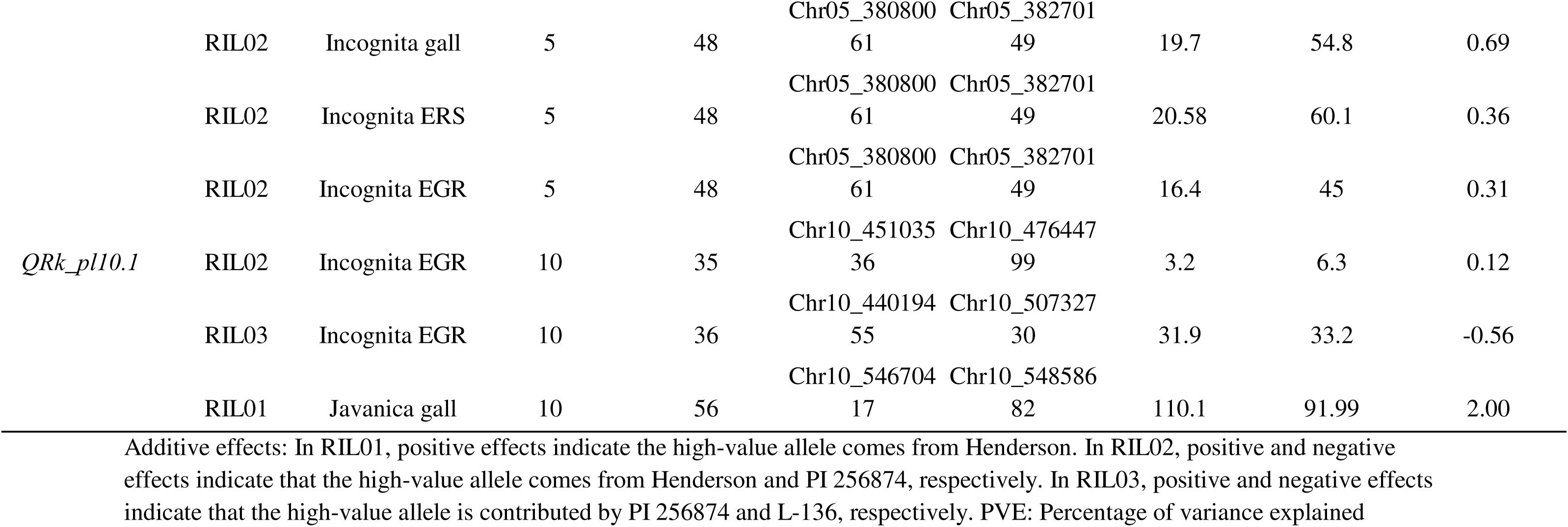
Chromosomal locations associated with root galling and reproduction response to different root-knot nematode species measured in 3 RIL populations Henderson x L-136 (RIL01), Henderson x PI 256874 (RIL02) and PI 256874 x L-136 (RIL03).

