## Supplementary Files for "Novel quantitative trait loci conferring broad-based resistance to root-knot nematodes in lima bean (*Phaseolus lunatus*)"

**Supplementary Figure 1:** RIL01, RIL02 and RIL03 Whole Genome LOD Peaks

(a.) depicts a whole genome representation of the LOD peaks associated with respective traits in the RIL01 population. (b.) depicts a whole genome representation of the LOD peaks associated with respective traits in the RIL02 population. Finally, (c.) depicts a whole genome representation of LOD peaks associated with respective traits in the RIL03 population.

a.

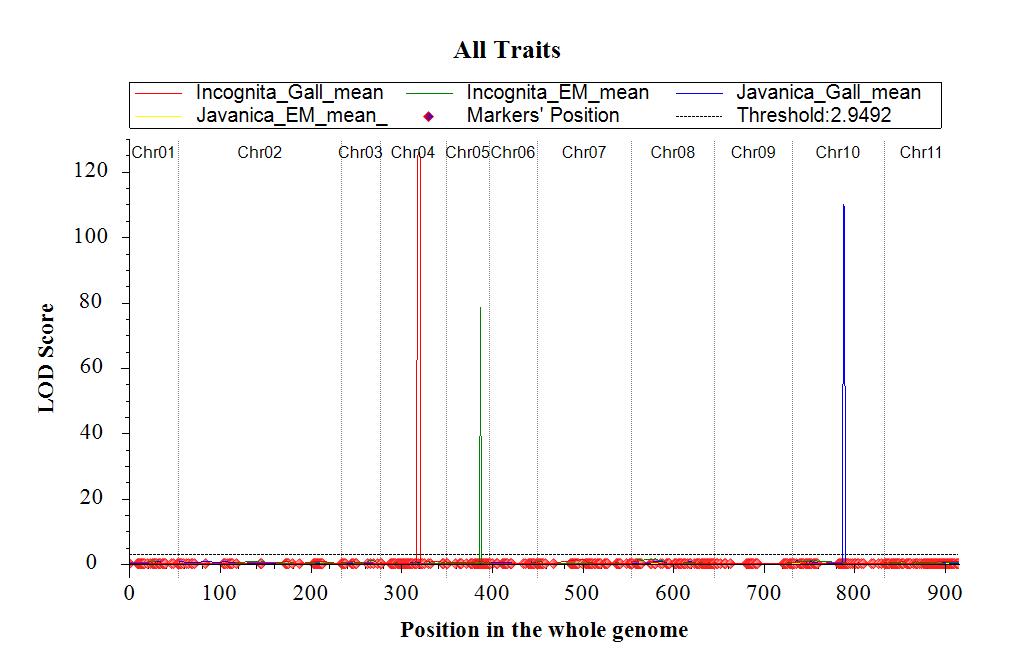

b.

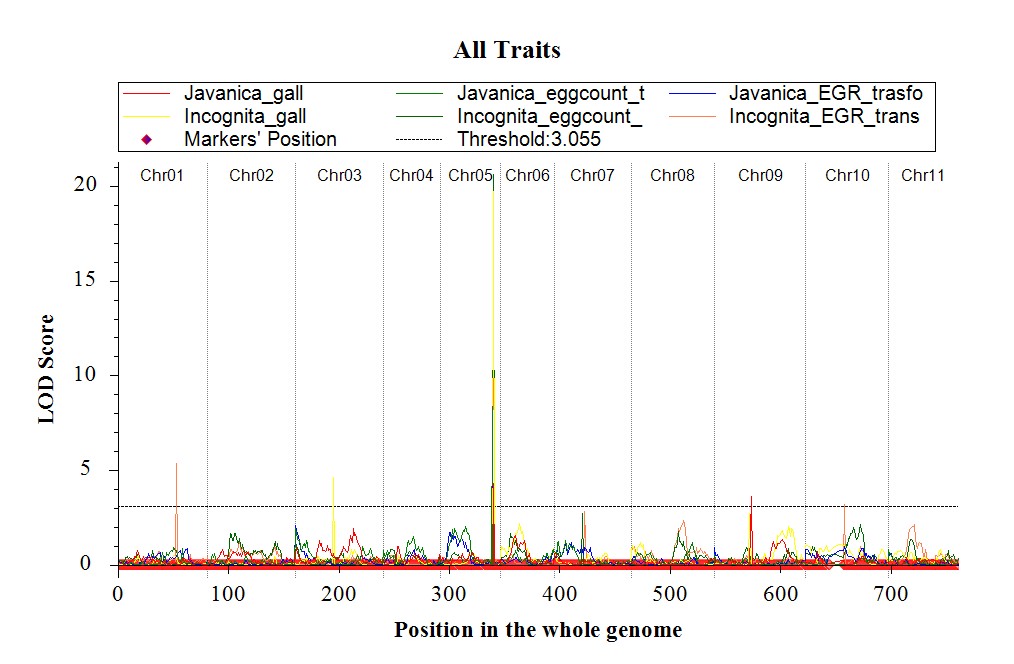

c.

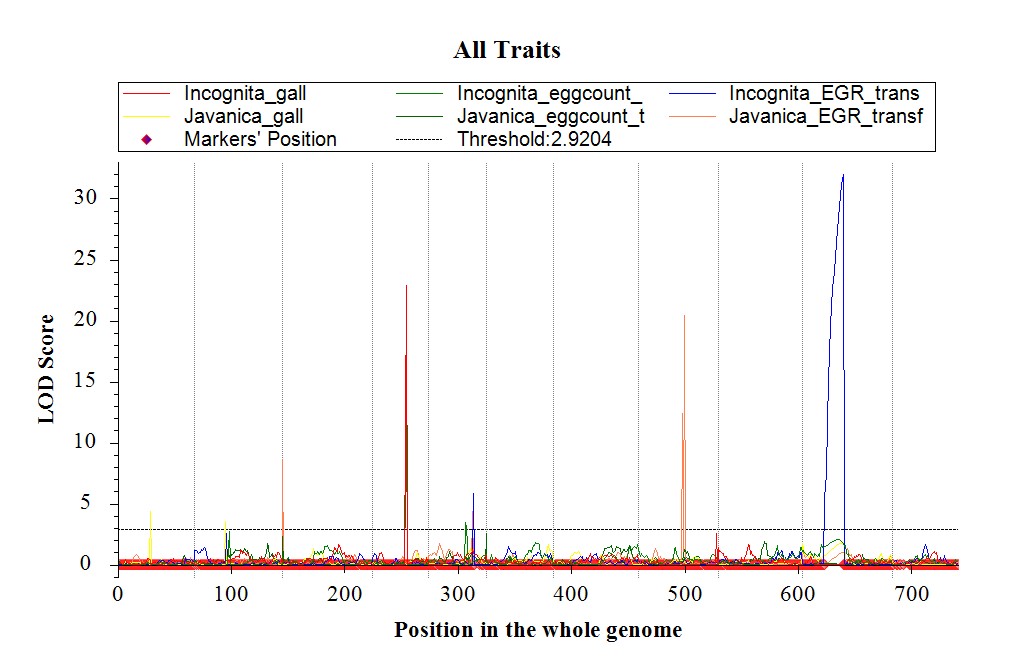

**Supplemental Table 1:** Data Table for RIL02 (Henderson x PI 256874) population

*Parental Phenotypes*

|  | Incognita total eggs (mean) | Incognita Egg Root Ratio (mean) | Javanica total eggs (mean) | Javanica Egg Root Ratio (mean) | Incognita Gall Mean | Javanica Gall Mean |
| --- | --- | --- | --- | --- | --- | --- |
| Henderson | 79,667 | 5,610 | 89,167 | 3,417 | 3.5 | 3.5 |
| PI 256874 | 10,667 | 175 | 22,000 | 430 | 1.35 | 0.6 |

*Phenotype Class 1 (lines with the PI phenotype) - 38 lines*

|  | Incognita total eggs | Incognita Egg Root Ratio | Javanica total eggs | Javanica Egg Root Ratio | Incognita Gall Mean | Javanica Gall Mean |
| --- | --- | --- | --- | --- | --- | --- |
| Mean | 22,250 | 754 | 23,556 | 808 | 1.555 | 1.1035 |
| Standard Deviation | 19,854 | 664 | 20,019 | 727 |  |  |

*Phenotype Class 2 (lines with Henderson phenotype) - 42 lines*

|  | Incognita total eggs | Incognita Egg Root Ratio | Javanica total eggs | Javanica Egg Root Ratio | Incognita Gall Mean | Javanica Gall Mean |
| --- | --- | --- | --- | --- | --- | --- |
| Mean | 146,453 | 4,303 | 137,313 | 4,082 | 3.1295 | 2.235 |
| Standard Deviation | 60,792 | 2,467 | 112,199 | 3,139 |  |  |

*Phenotype class 3: (low javanica galling) - 17 lines*

|  | Incognita total eggs | Incognita Egg Root Ratio | Javanica total eggs | Javanica Egg Root Ratio | Incognita Gall Mean | Javanica Gall Mean |
| --- | --- | --- | --- | --- | --- | --- |
| Mean | 106,559 | 5,701 | 56,794 | 3,481 | 2.5295 | 0.899 |
| Standard Deviation | 62,006 | 2,622 | 64,045 | 3,430 |  |  |

*Phenotype Class 4 (incognita resistance - low galling and low egg counts) - 13 lines*

|  | Incognita total eggs | Incognita Egg Root Ratio | Javanica total eggs | Javanica Egg Root Ratio | Incognita Gall Mean | Javanica Gall Mean |
| --- | --- | --- | --- | --- | --- | --- |
| Mean | 29,128 | 840 | 83,295 | 2,553 | 1.647 | 1.498 |
| Standard Deviation | 15,241 | 501 | 59,260 | 1,309 |  |  |

**Supplemental Table 2:** QTL validation table for RIL03 (PI256874 x L-136)

| Line | Donor parent @  *QRk_pl04.1* | Donor parent @ *QRk_pl05.1* | Donor parent @  *QRk_pl10.1* | Has putative resistance QTL? | Incognita Mean Galling | Incognita Mean Egg Count | Incognita Mean Egg Root Ratio | Javanica Mean Galling | Javanica Mean Egg Count | Javanica Mean Egg Root Ratio |
| --- | --- | --- | --- | --- | --- | --- | --- | --- | --- | --- |
| RIL047 | L-136 | PI 256874 | PI 256874 | yes | 4.67 | 6,482 | 25 | 1 | 401,111 | 1,965 |
| RIL063 | L-136 | PI 256874 | PI 256874 | yes | 1.83 | 13,889 | 67 | 0.5 | 111,389 | 1,632 |
| RIL070 | L-136 | PI 256874 | PI 256874 | yes | 0.83 | 236,111 | 710 | 0 | 33,333 | 147 |
| RIL072 | L-136 | PI 256874 | PI 256874 | yes | 3.50 | 42,593 | 170 | 2 | 356,667 | 1,954 |
| RIL081 | L-136 | PI 256874 | PI 256874 | yes | 3.50 | 927 | 4 | 0.5 | 283,333 | 1,691 |
| RIL083 | L-136 | PI 256874 | PI 256874 | yes | 3.50 | 103,704 | 266 | 1.5 | 628,889 | 2,556 |
| RIL016 | PI 256874 | L-136 | L-136 | no | 5.67 | 395,370 | 2,458 | 1 | 111,667 | 1,524 |
| RIL071 | PI 256874 | L-136 | L-136 | no | 7.00 | 505,555 | 3,621 | 3.5 | 2,026,667 | 29,067 |
| RIL087 | PI 256874 | L-136 | L-136 | no | 8.00 | 637,963 | 3,718 | 2.5 | 733,334 | 8,975 |
| PI 256874 | PI 256874 | PI 256874 | PI 256874 | partial | 5.92 | 404,167 | 1,628 | 2 | 315,370 | 1,594 |
| L-136 | L-136 | L-136 | L-136 | partial | 2.83 | 323,148 | 2,204 | 0 | 494,074 | 2,691 |
| Henderson | Henderson | Henderson | Henderson | no | 9.58 | 750,926 | 4,614 | 4 | 2,100,000 | 18,722 |

**Supplemental Table 3:** F1 phenotype tables for RIL01 (Henderson x L-136), RIL02 (Henderson x PIc256874) and RIL03 (PI 256874 x L-136) populations.

|  | Total Egg Count *Incognita* | Egg Count : Root Mass Ratio *Incognita* | Galling *Incognita* | Total Egg Count *Javanica* | Egg Count: Root Mass Ratio *Javanica* | Galling *Javanica* |
| --- | --- | --- | --- | --- | --- | --- |
| Henderson | 563,730 | 5,712 | + | 635,079 | 5,926 | + |
| L-136 | 162,143 | 1,343 | - | 449,603 | 3,431 | - |
| PI 256874 | 42,619 | 161 | - | 25,238 | 83 | - |
| Henderson x L-136 (F1) | 710,159 | 6,177 | - | 427,778 | 2,895 | - |
| Henderson x PI 256874 (F1) | 711,667 | 7,005 | + | 611,574 | 6,460 | + |

|  | Total Egg Count *Incognita* | Egg count: Root Mass Ratio *Incognita* | Galling *Incognita* | Total Egg Count *Javanica* | Egg Count: Root Mass Ratio *Javanica* | Galling *Javanica* |
| --- | --- | --- | --- | --- | --- | --- |
| Henderson | 426,667 | 3,381 | + | 339,722 | 3,629 | + |
| L-136 | 403,111 | 2,220 | - | 1,011,111 | 5,257 | - |
| PI 256874 | 31,111 | 104 | - | 78,556 | 221 | - |
| PI 256874 x L-136 (F1) | 524,723 | 2,000 | - | 4,422,444 | 22,638 | - |
